# Identification of N-linked glycans as specific mediators of neuronal uptake of acetylated α-Synuclein

**DOI:** 10.1101/407247

**Authors:** Melissa Birol, Slawomir P. Wojcik, Andrew D. Miranker, Elizabeth Rhoades

## Abstract

Cell-to-cell transmission of toxic forms of α-Synuclein (αS) is thought to underlie disease progression in Parkinson’s disease. αS in humans is constitutively N-terminally acetylated (αS_acetyl_), although the impact of this modification is relatively unexplored. Here we report that αS_acetyl_ is more effective at inducing intracellular aggregation in primary neurons than unmodified αS (αS_un_). We identify complex N-linked glycans as binding partners for αS_acetyl_, and demonstrate that cellular internalization of αS_acetyl_ is reduced significantly upon cleavage of extracellular N-linked glycans, but not other carbohydrates. We verify binding of αS_acetyl_ to N-linked glycans *in vitro*, using both isolated glycans and cell-derived proteoliposomes. Finally, we identify neurexin 1β, a neuronal glycoprotein, as capable of driving glycan-dependent uptake of αS_acetyl_. Importantly, our results are specific to αS_acetyl_ as αS_un_ does not demonstrate sensitivity for N-linked glycans. Our study identifies extracellular N-linked glycans, and neurexin 1β specifically, as key modulators of neuronal uptake of physiological αS_acetyl_ drawing attention to the potential therapeutic value of αS_acetyl_-glycan interactions.

## Introduction

The pathologies of Parkinson’s disease and related synucleinopathies are characterized by the accumulation of aggregates of the neuronal protein α-Synuclein (αS) (Goedert, 2001). The prevailing hypothesis is that toxicity is mediated by prefibrillar oligomers of αS (Luk et al, 2009). Emerging evidence suggests that cell-to-cell transmission of toxic αS species may be the basis of disease propagation (Guo & Lee, 2014).

αS is a small, soluble protein that is intrinsically disordered in the cytoplasm (Theillet et al, 2016). It binds peripherally to anionic lipid bilayers through its N-terminal domain, which becomes α-helical upon binding (Dikiy & Eliezer, 2014). The localization of αS to nerve terminals (Diao et al, 2013; George et al, 1995) and to cellular lipid raft domains (Fortin et al, 2004) suggests there are components or properties of cellular membranes which may be important for αS binding and function which may not be fully reproduced by simple lipid mixtures. Indeed, specific components of the extracellular membrane, including proteins (Mao et al, 2016) and proteoglycans (Ihse et al, 2017), have been identified as having roles in the uptake of pathogenic αS species.

αS is subject to various post-translational modifications, including phosphorylation, ubiquitination, glycation, acetylation and arginylation, some of which are correlated with pathology (Beyer & Ariza, 2013; de Oliveira et al, 2017; Vicente Miranda et al, 2017). Mass spectrometry analysis indicates that the majority of these modifications are found on only a very small fraction of αS (Kellie et al, 2014). N-terminal acetylation, however, is unique in that it is ubiquitously present on αS *in vivo* (Anderson et al, 2006; de Oliveira et al, 2017), in both healthy persons and Parkinson’s disease patients (Anderson et al, 2006; Kellie et al, 2014). In contrast to other protein modifications, including acetylation of side chains, acetylation of the amino terminus occurs co-translationally and is irreversible (Starheim et al, 2012). For many proteins, it is required for recognition of cellular binding partners (Scott et al, 2011). Work from our group (Trexler & Rhoades, 2012) and others have demonstrated that N-terminal acetylation alters the fundamental biophysical properties of αS; it moderately effects its binding to synthetic lipid bilayers (Dikiy & Eliezer, 2014; Kang et al, 2012) and rates of aggregation (Kang et al, 2012). How this modification impacts interactions with other cellular binding partners, and in particular the plasma membrane proteins which have been identified as receptors involved in cellular uptake and aggregation, has not been explored. Here, we investigate the role of N-terminal acetylation of αS on cellular internalization and aggregate propagation. Using cell biological and biophysical approaches, we demonstrate that N-terminal acetylation of αS confers interactions with N-linked glycans relevant to cellular uptake, identify neurexin 1β as a receptor for glycan-dependent uptake of αS and provide insight into the mechanism of cellular recognition relevant to uptake.

## Results

### N-terminal acetylation of αS enhances formation of intracellular aggregates in neurons

Recently observations that exogenously added aggregates of αS are capable of seeding aggregation of endogenous αS has prompted an interest in understanding the molecular players involved. However, investigations of this phenomenon have relied on aggregates comprised of unmodified αS (αS_un_). To determine whether N-terminal acetylation of αS (αS_acetyl_) alters this seeding behavior, primary hippocampal neurons were incubated with pre-formed fibrils (PFFs) of αS_acetyl_ or αS_un_ (Volpicelli-Daley et al, 2014) (Fig EV1). While both αS_acetyl_ and αS_un_ PFFs resulted in the formation of abundant intracellular aggregates (Fig 1A), the kinetics of aggregate formation differ significantly. For αS_un_ PFFs, the time-course was in good agreement with previously published studies (Volpicelli-Daley et al, 2014); aggregates were observed in axons by day 7 and had spread to somatodendritic compartments by day 10. For αS_acetyl_ PFFs, aggregates in axons were already prevalent by day 3 and spreading to somatodentric compartments was readily apparent by day 7. Measurements beyond 7 days were not possible for αS_acetyl_ PFFs due to significant cell death.

**Figure 1.**
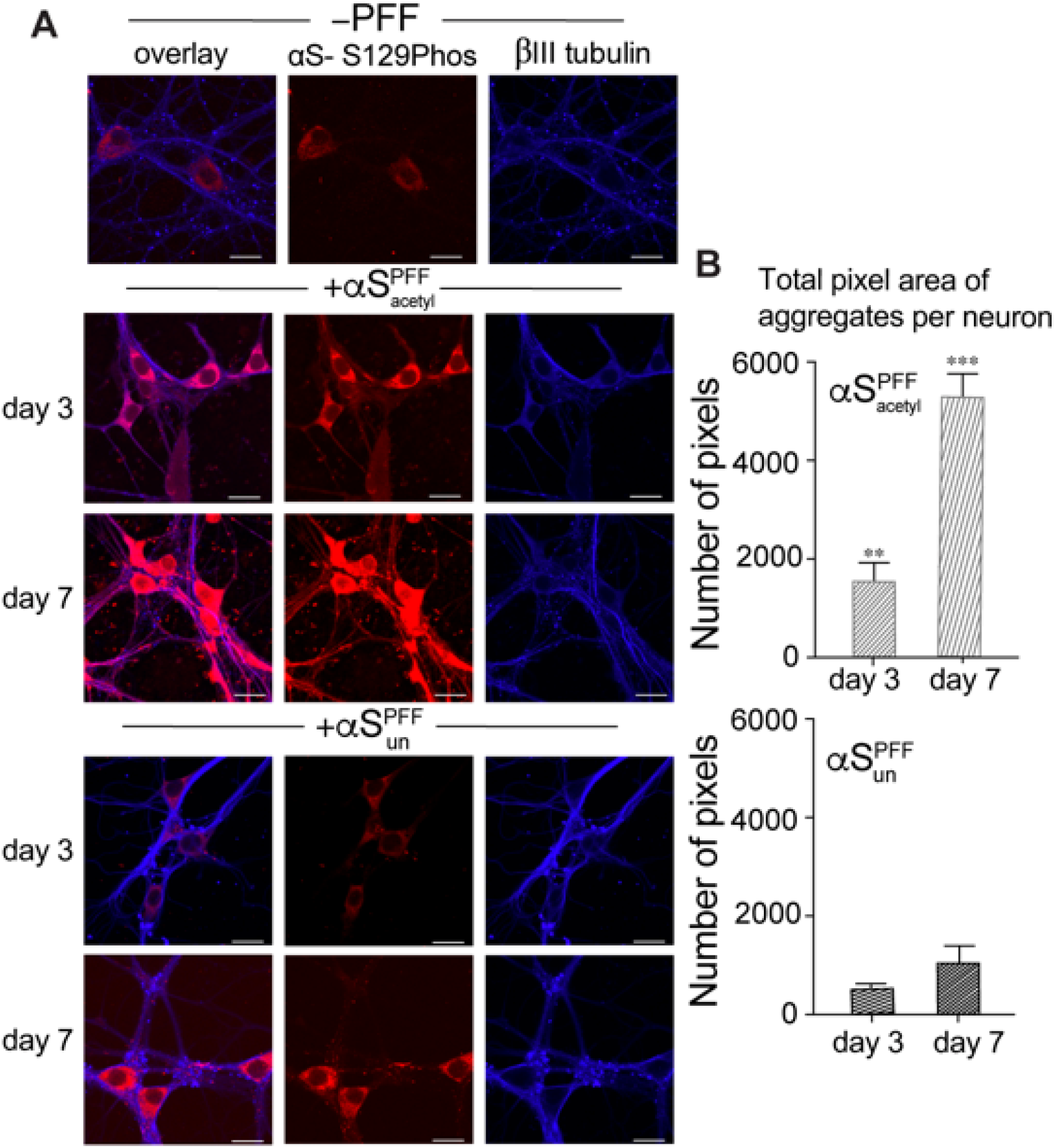
αS_acetyl_ PFFs are more effective at inducing pathological aggregates in primary neurons. A Representative images of aggregates of endogenous αS in cultured mouse hippocampal neurons following incubation with αS_acetyl_ or αS_un_ PFFs. Images shown after 3 and 7 days following treatment with PFFs. Red: αS-pS129 antibody; Blue: βIII-tubulin antibody. B Quantification of αS aggregates formed from images in (A). Aggregates in neurons treated with αS_acetyl_ PFFs are larger and more numerous. Data information: n=80 neurons, 3 independent experiments, *P<0.01, **P<0.001 and ***P<0.0001 by the Student’s T-Test compared to αS_un_ PFF treated neurons. The data are presented as mean±SD, n=3. Scale bars=20 μm.

To compare the effectiveness of αS_acetyl_ and αS_un_ PFFs in nucleating intracellular aggregate formation and growth, we quantified the total number of aggregates per neuron, reflecting the seeding capacity of the added αS PFFs (i.e., their ability to induce the initial formation of intracellular αS aggregates), as well as total aggregate area per neuron, reflecting the overall ability of the added αS PFFs to accelerate further aggregate growth. This quantification revealed that overall rate of aggregate formation is >2-fold faster for neurons treated with αS_acetyl_ PFFs as compared to αS_un_ PFFs (Fig 1B). This result demonstrates that αS_acetyl_ PFFs are markedly more potent seeds for pathological propagation of αS aggregation in neurons.

### αS_acetyl_ is more rapidly internalized by SH-SY5Y cells than αS_un_

This observation sparked our interest in determining the origin in differences in aggregate propagation in neurons. To do so, SH-SY5Y cells were chosen because they retain many of the pathways dysregulated in Parkinson’s disease and thus are widely used as a cellular model for disease. SH-SY5Y cells were incubated with monomer or PFF αS fluorescently labeled with Alexa Fluor 488 (αS-AL488). After 12 hours of incubation, monomer and PFF αS_acetyl_ and αS_un_ appeared as puncta, co-localized with an endosomal marker (Fig 2A, 2B, EV2A and EV2B). In order to make a more quantitative comparison of both kinetics and quantity of uptake between αS_acetyl_ and αS_un_, cellular internalization was measured as a function of time. Three orthogonal methods were used to quantify uptake: (1) loss of monomer αS from the cell media was measured using fluorescence correlation spectroscopy (FCS) (LaRochelle et al, 2015); (2) the amount of internalized monomer and PFF αS were quantified by confocal imaging; or (3) both extracellular and internalized monomer and PFF were quantified by polyacrylamide gel electrophoresis (Fig EV3). Results of all three of these approaches are consistent and reveal that both monomer and PFF αS_acetyl_ are internalized more rapidly (Fig 2C) and to a greater extent (Fig 2D and EV3) than αS_un_. Uptake was inhibited at 4°C, indicating that active endocytotic pathways are required (Fig EV4A). Control measurements made using non-neuronal lineage HEK cells found no internalization of either αS_acetyl_ or αS_un_ (Fig EV4B). Both SH-SY5Y and HEK cells showed rapid uptake of transferrin, indicating that lack of αS internalization is not due to inherent differences in rates of clathrin-dependent endocytosis between the cell lines (Fig EV4C).

**Figure 2.**
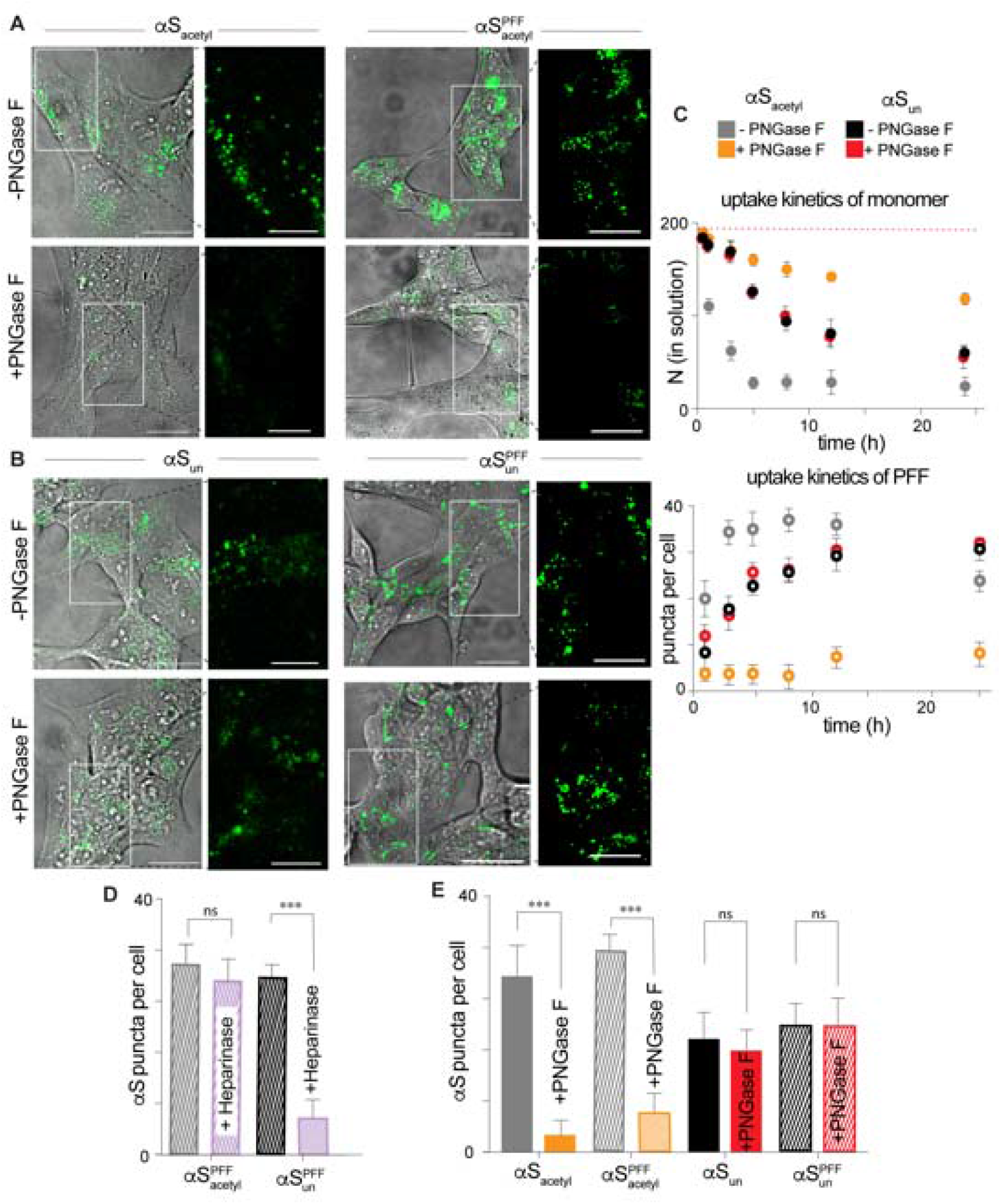
Complex N-linked glycans selectively enhance uptake of αS_acetyl_ by SH-SY5Y cells. A Representative images of SH-SY5Y cells following 12h incubation with monomer or PFF αS_acetyl_-AL488, -/+ PNGase F treatment. B As in (A) but for monomer or PFF αS_un_-AL488. C Upper: Kinetics of internalization by SH-SY5Y cells of monomer αS as quantified by loss from extracellular medium by FCS, -/+ PNGase F treatment. Lower: Kinetics of internalization by SH-SY5Y cells of PFF αS as quantified by puncta analysis of confocal images, -/+ PNGase F treatment. D Quantification of internalization of αS PFFs by SH-SY5Y cells, -/+ Heparinase I/III treatment. Images collected following 12 h incubation with protein and quantified by puncta analysis. E Quantification of internalization of αS monomer and PFF by SH-SY5Y cells, -/+ PNGase F treatment. Images collected following 12 h incubation with protein and quantified by puncta analysis. Data information: All protein uptake measurements with 200 nM monomer or 200 nM PFFs (monomer units, 20:1 αS:αS-AL488); n=100 cells, 3 independent experiments, *P<0.01, **P<0.001 and ***P<0.0001 by the Student’s T-Test). Scale bars=20 μm.

### Cleavage of extracellular N-linked glycans inhibits uptake of αS_acetyl_ by SH-SY5Y cells

Cell surface heparan sulfate proteoglycans have been observed to facilitate uptake of a number of fibrillar amyloid proteins, including αS_un_ (Holmes et al, 2013). To investigate the relevance of proteoglycans to αS_acetyl_ uptake, SH-SY5Y cells were treated with Heparinase I/III, an enzyme that cleaves these carbohydrates, for 6 hours. After exchange of media to remove the enzyme, αS-AL488 was added and incubated an additional 12 hours. This incubation period was chosen because it allows for reproducible quantification of puncta, and there is no evidence of protein or fluorophore degradation that may occur longer time points (Fig EV2B). In agreement with previous reports, we found that treatment of SH-SY5Y cells with Heparinase reduced the uptake of αS_un_ PFFs (Fig 2D and EV5A). Interestingly, however, Heparinase pre-treatment did not alter uptake of αS_acetyl_ PFFs nor that of αS_acetyl_ monomer (Fig 2D, EV5A and EV5B).

These results prompted us to consider other endoglycosidases, as the majority of cell-surface proteins are post-translationally modified by glycosylation (Moremen et al, 2012), including a number of proteins that have been identified as receptors for αS_un_ PFFs (Mao et al, 2016; Shrivastava et al, 2015; Urrea et al, 2017). Complex N-linked glycans were selectively removed from SH-SY5Y cells using peptide-N-glycosidase F (PNGase F). Following incubation with αS_acetyl_, a decrease in the number of intracellular puncta in the PNGase F treated cells relative to untreated cells was observed, both for monomer and PFF αS_acetyl_, reflecting a significant decrease in uptake by deglycosylated cells (Fig 2A, 2C, 2E and EV3). Removal of N-linked glycans was confirmed using concanavalin A (con A), a lectin which binds α-D-mannose and α-D-glucose moieties found on N-linked glycans (Fig EV5C). Control measurements showed that PNGase F treatment does not impact clathrin-dependent endocytosis (Fig EV5D). Under our measurement conditions (200 nM protein and 12 hours incubation), neither untreated nor PNGase F-treated cells show evidence of increased toxicity upon incubation with monomer or PFF αS_acetyl_ or αS_un_ (Fig EV5E). Moreover, and strikingly, internalization of monomer and PFF αS_un_ did not demonstrate any sensitivity to PNGase F treatment (Fig 2B, 2C and 2D). Lastly, we tested endoglycosidase H (Endo H), which cleaves high mannose N-linked carbohydrates and found only a minor impact with this enzyme on uptake of Sacetyl by SH-SY5Y cells (Fig EV5B).

Our observations in SH-SY5Y cells were corroborated in cultured primary hippocampal neurons. Monomer and PFF αS_acetyl_ and αS_un_ are readily internalized by primary hippocampal neurons (Fig 3A and 3B). Removal of extracellular N-linked glycans by PNGase F results in a 10-fold decrease in the amount of internalized αS_acetyl_ (Fig 3A), while no effect is observed for αS_un_ (Fig 3B). Similar to our measurements in SH-SY5Y cells, PNGase F treatment of neurons causes no defects in clathrin-dependent endocytosis (Fig EV4C).

**Figure 3.**
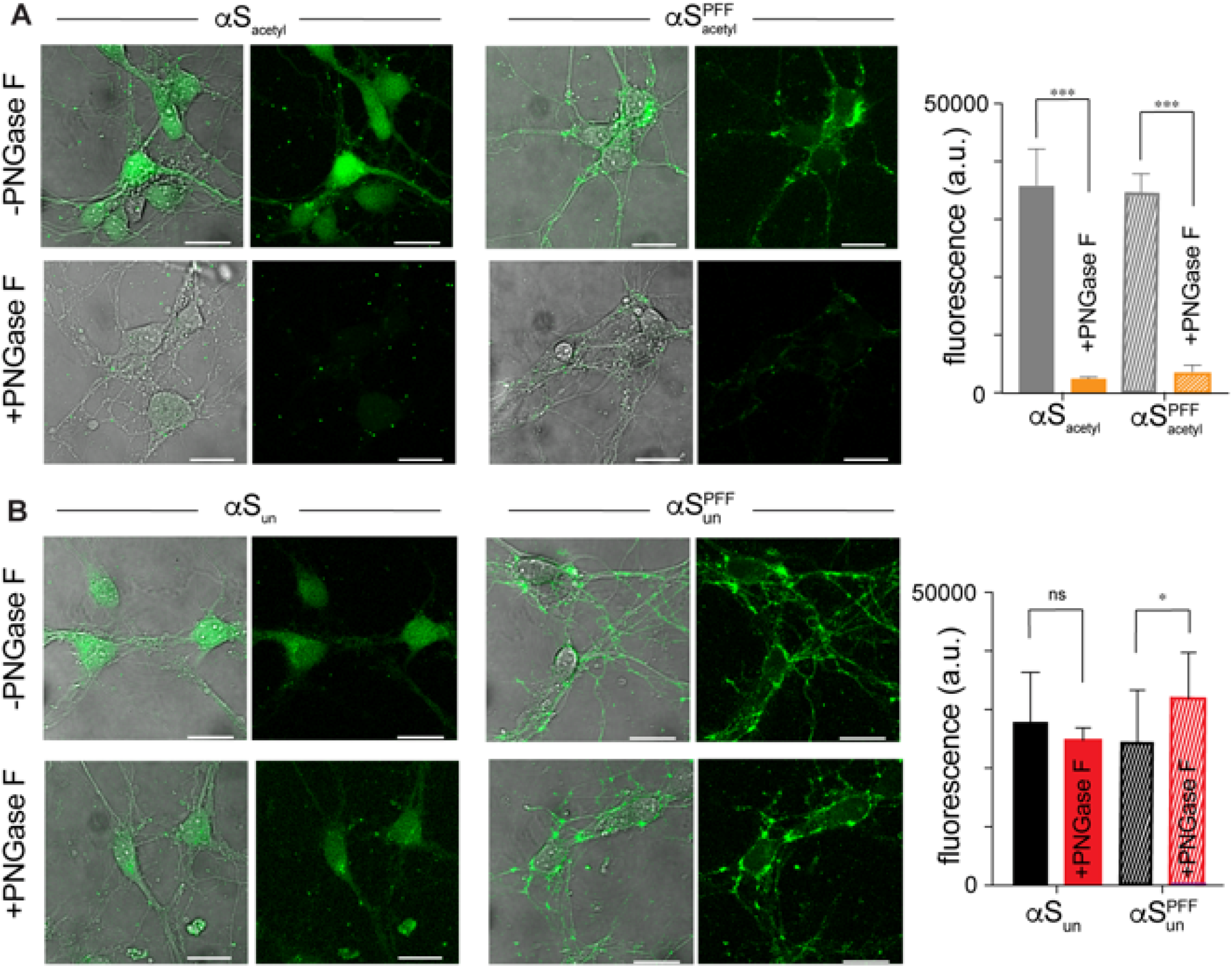
Complex N-linked glycans selectively enhance uptake of αS_acetyl_ by primary hippocampal neurons. A Representative images of mouse hippocampal neurons cells following 12h incubation with monomer or PFF αS_acetyl_-AL488, -/+ PNGase F treatment. Uptake quantified by total cellular fluorescence. B As in (A) but for monomer and PFF αS_un_-AL488. Data information: All internalization measurements with 200 nM monomer or 200 nM PFF (monomer units, 20:1 αS:αS-AL488); n=100 cells, 3 independent experiments, *P<0.01, **P<0.001 and ***P<0.0001 by the Student’s T-Test). Scale bars=20 μm.

### Removal of complex N-linked glycans alters αS binding to SH-SY5Y GPMVs

Our results thus far support specific interactions between αS_acetyl_ and complex, N-linked glycans found on neurons and SH-SY5Y, but not HEK, cells that drive internalization of both monomer and PFF αS. To investigate the molecular details of the interactions of αS with cell surface glycans, we used cell membrane-derived giant plasma membrane vesicles (GPMVs) which have a lipid and protein composition that closely resembles the cell plasma membrane (Bauer et al, 2009). Thus they serve as an excellent model of the cell membrane, but lack the active processes of cells, such as uptake, allowing for binding interactions to be observed. GPMVs were harvested from SH-SY5Y cells and incubated with monomer αS_acetyl_-AL488. αS_acetyl_ forms large, bright assemblies on the exterior of the SH-SY5Y GMPVs and causes their clustering, with larger assemblies and more clusters observed with increasing protein concentrations (Fig 4A and EV6A). Monomer αS_un_-AL488, on the other hand, binds uniformly (Fig 4B). The images present a striking contrast suggesting differences in binding affinity for αS_acetyl_ and αS_un_, which were quantified by FCS (Fig 4C). In these experiments, αS-AL488 was added GPMVs in a sample chamber and the amount of αS-AL488 that remained in solution after incubation was determined. After 60 minutes of incubation, 47±4% of αS_un_ and 80±6% of αS_acetyl_ were bound to the GPMVs (Fig 4C).

**Figure 4.**
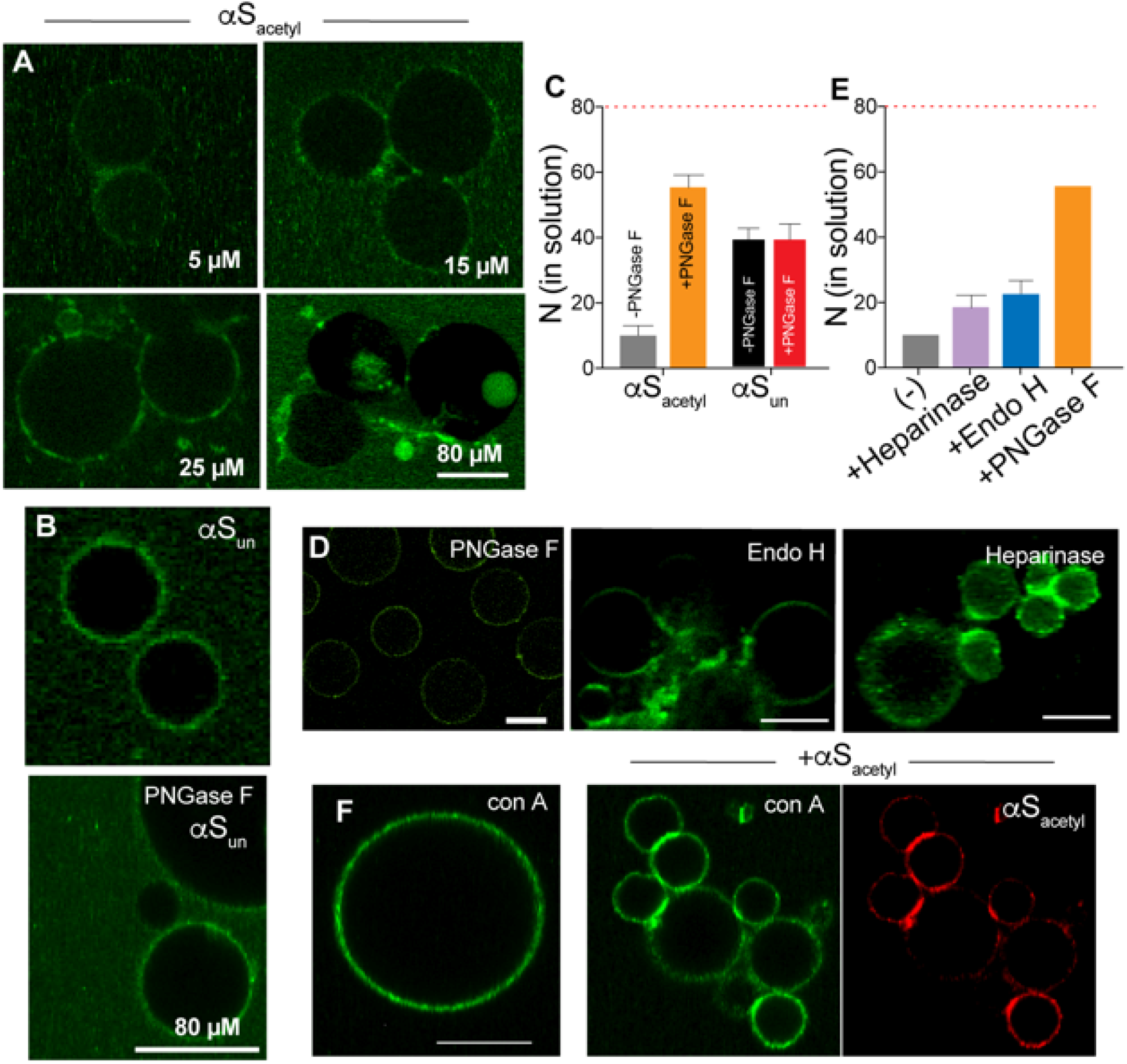
Removal of N-linked glycans disrupts binding of αS_acetyl_ to SH-SY5Y proteoliposomes.

A Images of SH-SY5Y GPMVs incubated with 100 nM αS_acetyl_-AL488 and unlabeled αS_acetyl_ (concentrations indicated).

B Upper: As in (A) but with 100 nM αS_un_-AL488 and 80 μM unlabeled αS_un_. Lower: As in upper panel but with treatment with PNGase F.

C Binding of 80 nM αS_acetyl_-AL488 or αS_un_-AL488 to GPMVs quantified by FCS as loss of protein (N, number of molecules) from solution. The number of molecules in control wells lacking GPMVs is indicated by the red-dashed line.

D Representative images of SH-SY5Y GPMVs incubated with 100 nM αS_acetyl_-AL488 and 80 μM unlabeled αS_acetyl_ after treatment of GPMVs with the indicated endoglycosidase.

E Binding of αS_acetyl_-AL488 to Heparinase- and Endo H-treated SH-SY5Y GPMVs measured by FCS, as in panel C. Binding +/- PNGase F from panel C shown for comparison.

F GPMVs incubated with 50 nM conA-AL488 in the absence and presence of 100 nM αS_acetyl_-AL594 and 80 μM of unlabeled αS_acetyl_.

Data information: The data are presented as mean±SD, n=3. Scale bars=20 μm.

Carbohydrates were selectively removed from the extracellular surface of the SH-SY5Y GPMVs by incubation with the same endoglycosidases previously used in our cell uptake measurements. Treatment with PNGase F results in a loss of inhomogenous binding and the bright αS_acetyl_ assemblies (Fig 4D and EV6B), as well as significantly decreasing the amount of bound αS_acetyl_ (Fig 4E). In contrast, binding of αS_un_ to SH-SY5Y GPMVs is unaltered by PNGase F treatment (Fig 4B). By comparison, bright αS_acetyl_ assemblies are still observed after treatment with Heparinase or Endo H (Fig 4D, 4E and EV6C), consistent with our cellular uptake measurements. Also consistent with our cellular uptake measurements is that only weak binding is observed for αS_acetyl_ to GPMVs derived from HEK cells (Fig EV6D).

Uniform binding of Alexa 488 labeled conA (conA-AL488) to SH-SY5Y GPMVs demonstrates that glycoproteins are distributed throughout the GPMV bilayer in the absence of αS_acetyl_ (Fig 4F and EV6E); the addition of αS_acetyl_ results in clustering of conA-stained proteins (Fig 4F, EV6F and EV6G). αS_acetyl_ appears to induce assembly through binding to and cross-linking multiple membrane glycoproteins, likely observed because GPMVs lack an intact cytoskeleton which would restrict large-scale rearrangement of plasma membrane proteins.

### αS_acetyl_ binds complex N-linked glycans with a distinct structure

Because its native function is thought to involve interactions with cellular membranes, binding of αS to synthetic lipid vesicles has been thoroughly investigated by a number of experimental methods (Eliezer et al, 2001; Kamp & Beyer, 2006; Rhoades et al, 2006). Membrane binding to synthetic lipid vesicles is mediated through the first ~95 residues of αS, which form an elongated α-helix upon association (Eliezer et al, 2001; Trexler & Rhoades, 2009). Intramolecular FRET measurements of αS bound to GPMVs were made; αS was labeled at residues 9 and 72, positions encompassing much of membrane-binding domain, which we have used previously as a probe of the elongated α-helical structure (Trexler & Rhoades, 2009). Mean energy transfer efficiencies (ET_eff_) of αS_acetyl_ and αS_un_ bound to SH-SY5Y GPMVs, were 0.43±0.08 and 0.21±0.06, respectively (Fig 5A and 5B). This lower ET_eff_ measured for αS_un_ is consistent with an elongated helix as measured by single molecule FRET with synthetic lipid bilayers (Trexler & Rhoades, 2009). The higher ET_eff_ measured for αS_acetyl_ demonstrates that it binds in a distinct conformation. Strikingly, when complex N-linked glycans are removed from the GPMV by treatment with PNGase F, ET_eff_ distributions peaked at 0.19±0.06 and 0.17±0.03 are observed for αS_acetyl_ and αS_un_, respectively (Fig 5A and 5B). Our interpretation of these findings is that in the absence of complex N-linked glycans, αS_acetyl_ binds to GPMVs through interactions with the lipid bilayer resulting in a predominantly extended conformation. In the presence of N-linked glycans, αS_acetyl_ binding is enhanced and it assumes as conformation distinct from the extended, membrane-bound α-helix. Binding of αS_un_, on the other hand, occurs through its interactions with the lipid bilayer resulting in the extended helical state even in the presence of N-linked glycans. These results provide a basis for understanding why no differences in uptake are observed binding (Fig 3C) or cellular uptake (Fig 2C and 2E) for αS_un_ upon PNGase F treatment.

**Figure 5.**
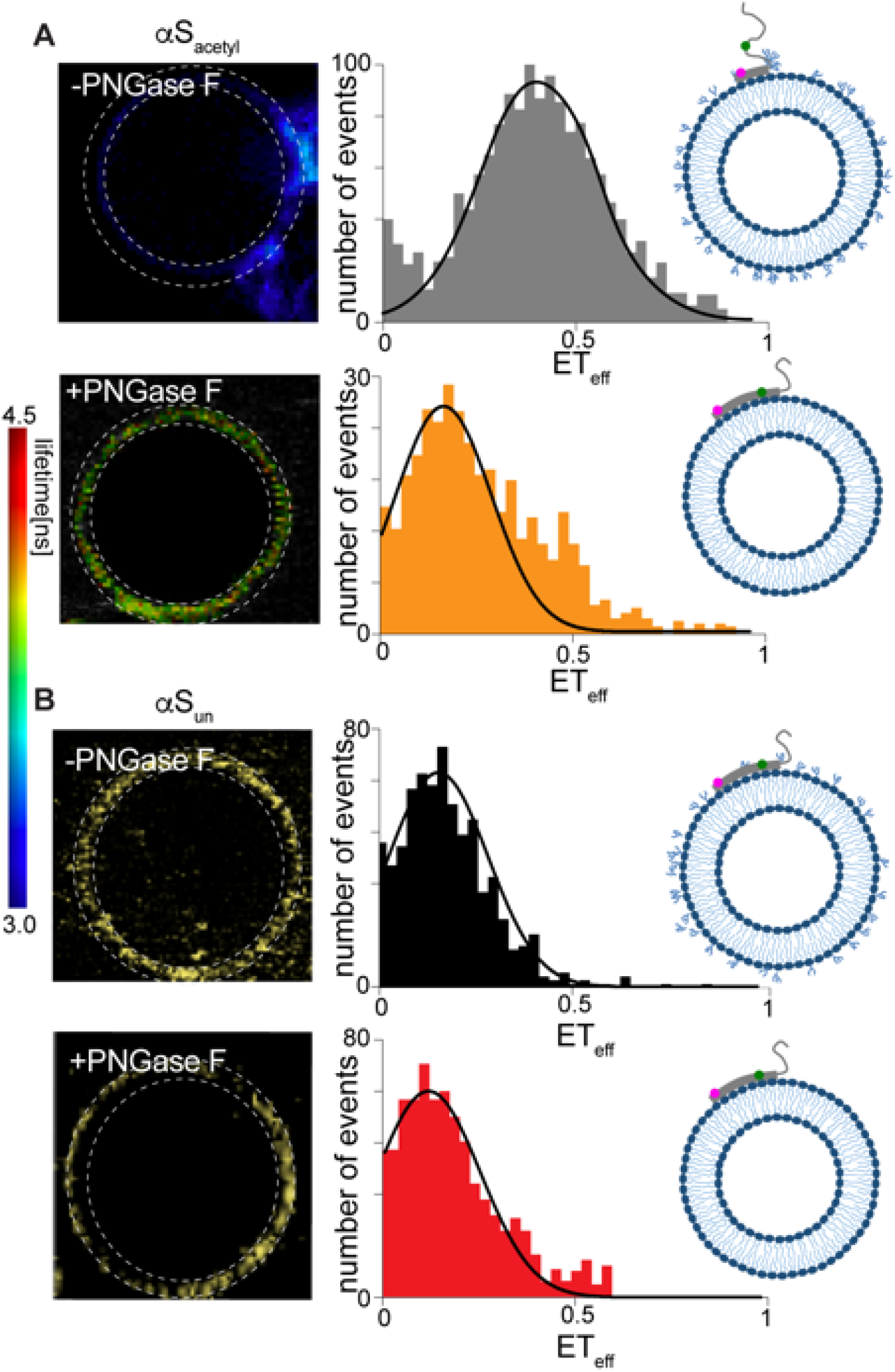
Intramolecular FLIM-FRET measurements of αS bound to SH-SY5Y GPMVs. A FLIM-FRET measurements of αS_acetyl_ bound to GPMVs without (upper) and with (lower) treatment with PNGase F Left: Representative image of a GPMV, colored by donor fluorophore lifetime shown in the scale bar. Right: Histogram of ET_eff_ calculated from the images, with Gaussian fits shown. The pixels used to calculate the histograms are indicated by dashed lines on the GPMV images. B As in (A) but with αS_un_. Data information: Histograms indicated are an average of three GPMVs per biological replicate (n=3).

### αS_acetyl_ binds isolated N-linked glycans from SH-SY5Y cells

To identify whether αS_acetyl_ binding to glycans requires either the associated glycoproteins or a lipid bilayer, binding of αS_acetyl_ to glycans in solution was measured by FCS. SH-SY5Y cells were treated with each of the three endoglycosidases used with the GPMVs, and the cleaved carbohydrates were isolated. The carbohydrates were titrated into αS_acetyl_-AL488 and the autocorrelation curves (Fig EV7A) were fit to extract the diffusion time and average number of fluorescent molecules, N (Fig 6A and EV7B). The diffusion time, which reflects the hydrodynamic size of the diffusing species, of αS_acetyl_ increases more than 25% with increasing concentrations of PNGase F-derived glycans, comparable to con A-AL488 (Fig 6A, 6B and EV7C). No increase in the diffusion time of αS_un_ is observed in the presence of PNGase F-derived glycans (Fig 6A). Similarly, the addition of carbohydrates obtained from Endo H or Heparinase treatment of SH-SY5Y cells or PNGase F treatment of HEK cells results in minimal changes in the diffusion time of αS_acetyl_ by FCS (Fig 6B and EV7D). NMR measurements show non-uniform glycan dependent changes in αS_acetyl_ peak intensity in the presence of PNGase F-cleaved glycans, but not simple carbohydrates (Fig 6C and EV7E). By a filtration-based assay, both monomer and PFF αS_acetyl_ are found to pull-down PNGase F-cleaved glycans from solution (Fig EV7F). In contrast to GPMV images, there is no evidence of glycan-mediated assembly of αS_acetyl_ in solution (Fig EV7C). This may reflect a difference in αS_acetyl_ binding to disperse glycans in solution compared to the relatively high density of glycans on the GPMV. Alternatively, it could be due to the absence of the relevant glycoprotein(s) as αS_acetyl_ binding may involve interactions both with N-linked glycans, as well as their associated glycoproteins.

**Figure 6.**
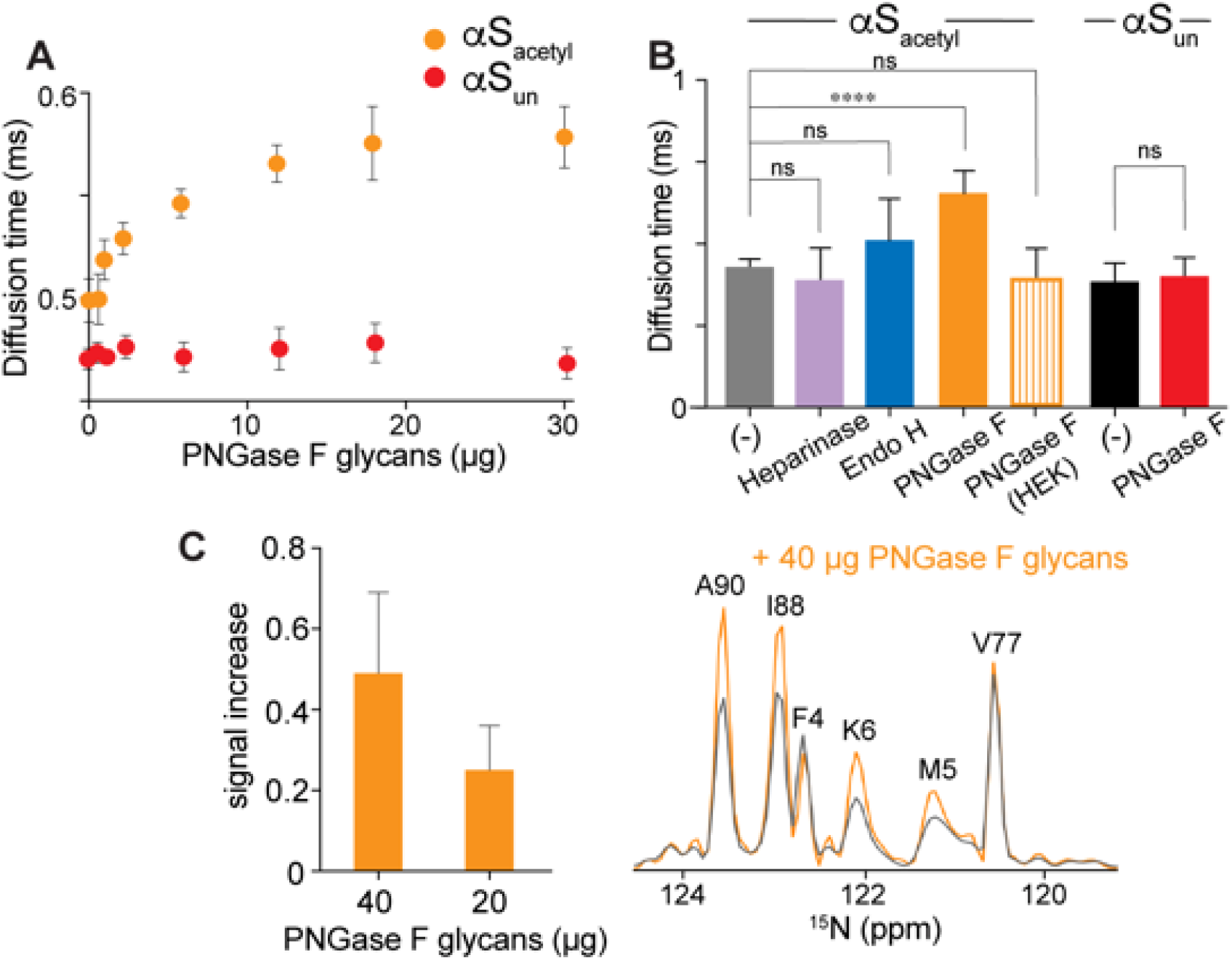
Binding of αS_acetyl_ to isolated cell surface N-linked glycans. A Diffusion time of αS_acetyl_-AL488 and αS_un_-AL488 (80 nM) as a function of PNGase F-derived glycan concentration. B Comparison of αS_acetyl_-AL488 and αS_un_-AL488 binding to carbohydrates (30 μg) cleaved by the indicated endoglycosidases. C (left) Percent increase in peak intensity for selected residues strongly enhanced by glycan binding and (right) cross-sections from ^15^N-^1^H HSQC spectra along the nitrogen dimension at 8.15 ppm for αS_acetyl_ in the absence (gray) and in the presence of PNGase F-derived glycans (orange). Data information: For (A-C), data are presented as mean±S.D., n=3. *P<0.01, **P<0.001 and ***P<0.0001 by the Student’s T-Test).

### Neurexin 1β drives internalization of αS_acetyl_

Our results to this point demonstrate that the uptake of both monomer and PFF αS_acetyl_ by SH-SY5Y cells or primary neurons is strongly impacted by its interactions with complex, N-linked glycans. Our biophysical measurements with GPMVs and isolated carbohydrates support this observation. This prompted us to try to identify a specific glycoprotein capable of binding to αS_acetyl_. One recent screen of transmembrane proteins identified neurexin 1β and lymphocyte activation gene 3 (LAG3), both of which contain N-linked glycosylation sites in their extracellular domains, as binding partners for αS_un_ (Mao et al, 2016). Although this study did not address the impact of N-terminal acetylation of αS, nor of glycosylation of the neurexin 1β or LAG3, it found that both receptor proteins exhibit specificity for PFF over monomer αS_un_, with the effect more striking for LAG3. To specifically address a possible role for glycosylation of these proteins in αS uptake, HEK cells were transfected with either LAG3 or neurexin 1β, each bearing a GFP tag on its intracellular domain (Fig EV8A). αS was labeled with Alexa Fluor 594 (αS-AL594) to allow for simultaneous imaging of the transfected protein and exogenously added αS. In the absence of LAG3 or neurexin 1β, no uptake of αS_acetyl_ or αS_un_ monomer or PFF by HEK cells was observed (Fig EV4B). At comparable transfection levels, the proteins localized to the plasma membrane, as expected (Fig EV8B). Upon the addition of either αS_acetyl_ or αS_un_ PFFs to LAG3 expressing cells, colocalization with LAG3 on the plasma membrane followed by uptake was observed (Fig 7A). Consistent with the results of the screen which identified these proteins, neither colocalization nor uptake were detected for monomer αS_un_ (Fig7C and EV8C), nor did we observe it for monomer αS_acetyl_ (Fig 7A and 7C). Treatment of the LAG3-expressing cells with PNGaseF did not decrease uptake of αS_un_ or αS_acetyl_ PFFs (Fig 7B and 7C).

**Figure 7.**
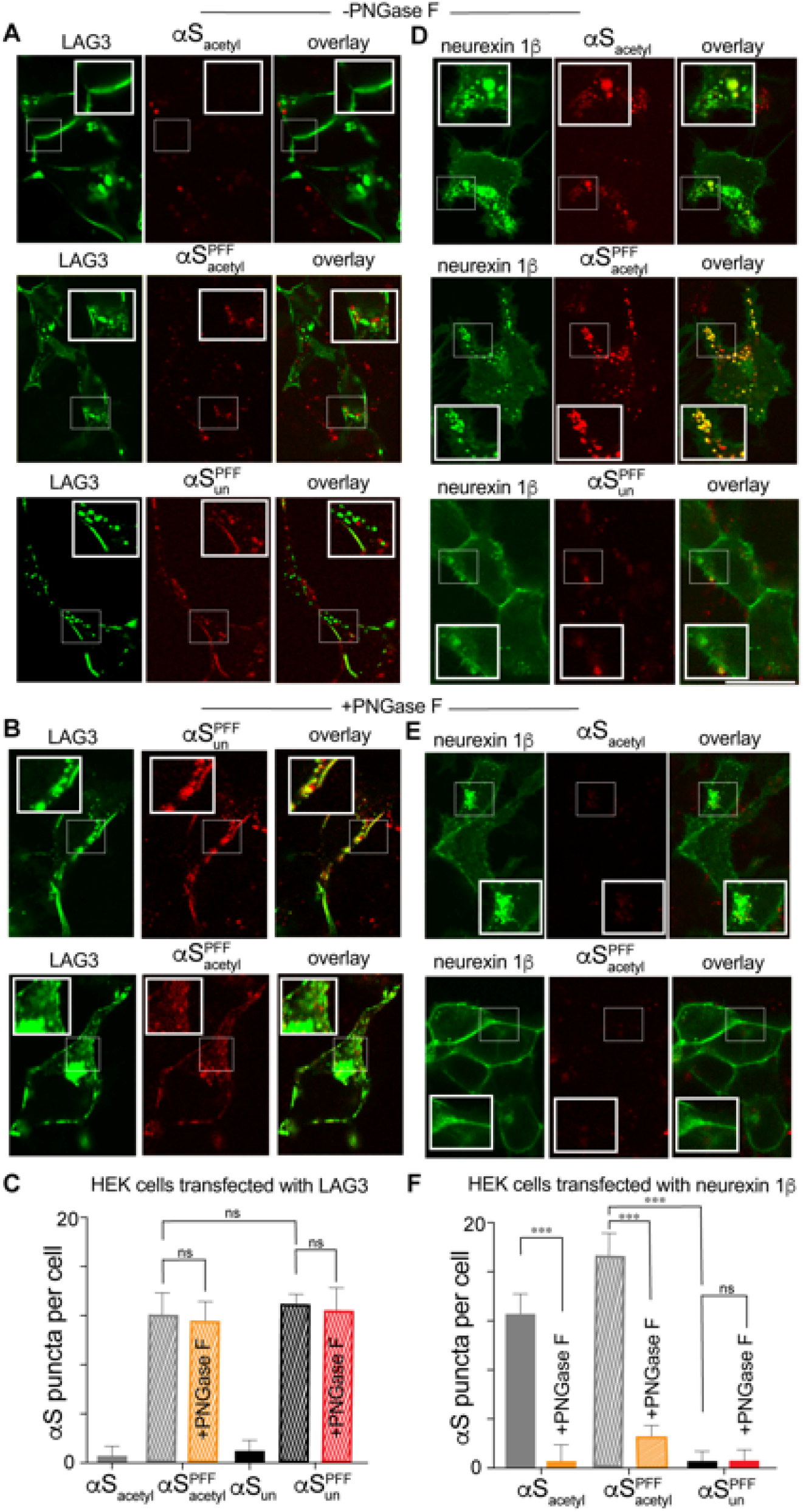
Glycosylated neurexin 1β is a receptor for αS_acetyl_. A HEK cells transfected with LAG3-GFP after incubation with monomer or PFF αS_acetyl_-AL594 or PFF αS_un_-AL594 B HEK cells transfected with LAG3-GFP but cells treated with PNGase F prior to incubation with monomer or PFF αS_acetyl_-AL594 or PFF αS_un_-AL594. C Quantification of uptake of monomer and PFF αS-AL594 by LAG3-GFP transfected HEK cells and +/- PNGase F treatment quantified by puncta analysis (for αS_acetyl_ and αS_un_ PFFs). D As in (A) but HEK cells transfected with neurexin 1β-GFP. E As in (B) but HEK cells transfected with neurexin 1β-GFP. F As in (C) but HEK cells transfected with neurexin 1β-GFP.

The results from the neurexin 1β expressing HEK cells stand in striking contrast. Both monomer and PFF αS_acetyl_ were internalized by these cells where they colocalized in distinct intracellular puncta (Fig 7D and 7F). Intracellular puncta containing neurexin 1β were not observed in the absence of αS (Fig EV8A), demonstrating that αS_acetyl_ drives internalization of neurexin 1β during its uptake. We saw no evidence of binding or uptake of monomer or PFF αS_un_ in neurexin 1β expressing HEK cells (Fig 7D, 7F and EV8D). Treatment of neurexin 1β transfected HEK cells with PNGase F greatly decreased binding and uptake of αS_acetyl_ monomer and PFF (Fig 7E and 7F). Moreover, neurexin 1β maintained its plasma membrane localization. These data identify neurexin 1β and its glycosylation as key modulators of pathogenic cell-to-cell transmission of αS_acetyl_.

## Discussion

In the present study, we present compelling evidence that cellular internalization of αS_acetyl_ by primary neurons and SH-SY5Y cells is dependent upon complex, N-linked glycans. αS_acetyl_ binds to these complex, N-linked glycans in solution and on cell-derived proteoliposomes. Glycan-binding is a novel result for αS_acetyl_ and we propose that it is likely to be critical for recognition of functional protein binding partners. We identify one of those binding partners, neurexin 1β, and show that both its binding to αS_acetyl_ and the consequent uptake of αS_acetyl_ are dependent upon its glycosylation.

Critically, the underlying factor in our discovery is our use of αS_acetyl_ and our ability to make direct comparisons of between αS_acetyl_ and αS_un_. Although it shown more than ten years ago that αS_acetyl_ is the physiological form of αS (Anderson et al, 2006) and αS_acetyl_ can be produced in e. coli (Trexler & Rhoades, 2012), many studies, including those aimed at understanding function and uptake mechanisms of αS, still rely on the unmodified protein. Up to 80% of mammalian proteins are modified by N-terminal acetylation and for many, e.g. tropomyosin binding to actin (Hitchcock-DeGregori & Heald, 1987), this modification is required for recognition of binding partners (Drazic et al, 2016). Our current work demonstrates that this is true for αS_acetyl_ and neurexin 1β (Fig 7) and may also be true for other cellular binding partners of αS. Moreover, the increased uptake (Fig 2 and 3), and the resulting enhancement of intracellular aggregate formation induced by αS_acetyl_ PFFs relative to αS_un_ PFFs (Fig 1) provide compelling evidence that N-terminal acetylation of αS has physiological consequences which thus far have been overlooked. We anticipate that our results may help reconcile conflicting cellular/animal models with biochemical/biophysical studies of αS where proteins may lack the appropriate modifications.

Disordered proteins such as αS_acetyl_ often participate in highly specific, but relatively low affinity, interactions(Wright & Dyson, 2015). As such, multivalency provides sufficient avidity for biological interactions between disordered proteins and binding partners *in vivo*, in examples as diverse as tubulin polymerization (Li & Rhoades, 2017), liquid-liquid phase separation (Li et al, 2012) and nuclear transport (Milles et al, 2015). Intriguingly, interactions between glycan binding proteins, including lectins, and their binding partners are often described in the same terms. The binding between these proteins and a single glycan is often relatively low affinity (μM-mM) (Varki et al, 2009). We estimate the apparent K_d_ for monomer αS_acetyl_ and PNGase F-derived glycans from the FCS data shown in Fig. 6a to be ~ 10-20 μM. Multivalency both enhances the affinity and confers specificity on the interaction, as high affinity binding only occurs when the correct cluster of glycans is present and in the correct orientation. This requirement may underlie the differences we observe for αS_acetyl_ in its interactions with SH-SY5Y and primary neurons relative to HEK cells. While complex N-linked glycans are abundant on all mammalian cells membranes, there are significant differences in the specific glycome between cell types (Park et al, 2018). Most glycan binding proteins are members of well-characterized families and share similar structures or amino acid sequences (Van Holle et al, 2017). To our knowledge, there are no prior examples of entirely disordered proteins showing selective binding to complex, N-linked glycans, further underscoring the novelty of our results.

Monomer αS is disordered in solution, with transient helical structure in the amino terminus (Eliezer et al, 2001). N-terminal acetylation enhances the helical propensity of the first 12 residues of αS, as well as exerting long-range effects up through approximately residue 50 (Kang et al, 2011). These changes in conformational sampling conferred by N-terminal acetylation may have a role in the selective glycan binding of αS_acetyl_ relative to αS_un_. Also relevant to our study, which finds that both monomer and PFF αS_acetyl_ interact with N-linked glycans, the first ~45 residues of αS remain flexible and extended – and thus presumably available for binding – in fibrillar aggregates (Tuttle et al, 2016). That both monomer and PFF αS_acetyl_ and αS_un_ are internalized by SH-SY5Y cells, while only αS_acetyl_ uptake is impacted by N-linked glycans and only PFF αS_un_ uptake is impacted by Heparinase strongly suggests that there are multiple modes by which αS interacts with the extracellular plasma membrane. For αS_acetyl_, binding to the extracellular membrane is driven by its specific interactions with N-linked glycans. For αS_un_, binding appears to be derived primarily from interactions with outer leaflet lipids and, for PFF αS_un_, proteoglycans. In all cases, altering the amount of bound αS, whether by cleaving a specific carbohydrate or increasing the amount of glycoprotein (as with neurexin 1β), correspondingly alters the amount of internalized αS.

One consequence of our identification of αS_acetyl_ as a glycan binding protein is that it elicits a reconsideration of interactions between αS and putative binding partners. Many of the proteins identified as such, including glucocerebrosidase (Yap et al, 2011) and Rab3b (Cooper et al, 2006), as well as the LAG3 and neurexin 1β (Mao et al, 2016) examined here, contain N-linked glycosylation sites. Consistent with the description of glycan binding proteins above, our results with LAG3 and neurexin 1β emphasize the relevance of the specific glycans modifying the glycoprotein in determining binding of αS_acetyl_. Both LAG3 and neurexin 1β are modified by N-linked glycans; however, only the glycans on neurexin 1β mediate binding and uptake of αS_acetyl_ (Fig 7). As αS is a major target for drug development to treat Parkinson’s disease and other synucleinopathies, identification of αS_acetyl_ as a glycan binding protein provides new considerations for therapeutic approaches.

## Material and Methods

### αS expression and purification

αS was expressed in *E. coli* BL21 cells; for N-terminally acetylated αS, BL21 stocks containing the NatB plasmid with orthogonal antibiotic resistance were used. The purification of both acetylated (αS_acetyl_) and unmodified (αS_un_) protein was carried out as previously described (Trexler & Rhoades, 2012), with minor modifications. Briefly, two ammonium sulfate cuts were used (0.116 g/mL and 0.244 g/mL) with αS precipitating in the second step. The pellet was resolubilized in Buffer A (25 mM Tris pH 8.0, 20 mM NaCl, 1 mM EDTA) with 1 mM PMSF and dialyzed against Buffer A to remove ammonium sulfate. Dialyzed samples were then loaded to an anion exchange column (GE HiTrap Q HP, 5ml) and eluted with a gradient to 1 M NaCl. αS elutes at approximately 300 mM NaCl. Fractions containing αS were pooled and concentrated using Amicon Ultra concentrators (3000 Da MWCO). Concentrated samples were then loaded to a size exclusion column (GE HiLoad 16/600 Superdex75) and eluted at 0.5 ml/min. Fractions containing αS were again pooled and concentrated, then stored at −80°C. All αS constructs used in this work were checked by MALDI to confirm correct mass and presence of acetylation (Fig EV1).

For NMR measurements, 15N-labeled αS was grown in E. coli BL21 stocks containing the NatB plasmid in minimal medium (6 g/L Na_2_HPO_4_.7H_2_O, 3 g/L KH_2_PO_4_, 0.5 g/L NaCl, 1mM MgSO_4_, 300 μM CaCl_2_, 0.5 g/L ^15^NH_4_Cl) instead of LB medium and purified as described above.

### αS labeling

αS was site-specifically labeled at a single position by introduction of a cysteine at either residue 9 or residue 130. For labeling reactions, freshly purified αS (typically 200–300 μL of ~200 μM protein) was incubated with 1 mM DTT for 30 min at room temperature to reduce the cysteine. The protein solution was passed over two coupled HiTrap Desalting Columns (GE Life Sciences) to remove DTT and buffer exchanged into 20 mM Tris pH 7.4, 50 mM NaCl, 6 M guanidine hydrochloride (GdmCl). The protein was incubated overnight at 4 °C with stirring with 4x molar excess Alexa Fluor 488 (AL488) or Alexa Fluor 594 (AL594) maleimide (Invitrogen). The labeled protein was concentrated and buffer exchanged into 20 mM Tris pH 7.4, 50 mM NaCl using an Amicon Ultra 3K Concentrator (Millipore), with final removal of unreacted dye and remaining GdmCl by passing again over a set of coupled desalting columns equilibrated with 20 mM Tris pH 7.4, 50 mM NaCl.

For dual fluorophore labeling for FRET measurements, cysteines were introduced at residues 9 and 72. The protein was labeled as described above, with the following modifications. The protein was first incubated with donor fluorophore AL488 maleimide at a ratio of protein:dye of 1:0.5 for 2 h at room temperature with stirring. Then, 4x molar excess of acceptor fluorophore AL594 maleimide (Invitrogen) was added, and the reaction was continued overnight at 4 °C. The labeled protein was separated from unreacted dye as described above. αS labeled at these positions have been extensively studied in our lab; as documented in our previous publications, they do not perturb αS binding to lipid membranes and serve as excellent reporters of different conformations of membrane-associated αS (Middleton & Rhoades, 2010; Trexler & Rhoades, 2009).

### Fibril Formation

αS pre-formed fibrils (PFFs) were prepared as previously described (Volpicelli-Daley et al, 2014). Briefly, 100 μM αS was mixed with 5 μM αS-AL488 in 20 mM Tris pH 7.4 and 100 mM NaCl. To induce aggregation, this solution was incubated at 37°C for 5 days with agitation (1500 rpm on an IKA MS3 digital orbital shaker) in parafilm-sealed 1.5 mL Eppendorf tubes to ensure minimal solvent evaporation. The aggregation reaction was analyzed by Congo Red absorbance by diluting 10 μl of the aggregation solution in 140 μl 20 μM Congo Red. The mature fibrils were then pelleted by centrifugation (13,200 rpm for 90 mins at 4°C) and the supernatant removed. Fibers were resuspended in an equal volume (relative to supernatant) of 20 mM Tris pH 7.4, 100 mM NaCl. Mature fibers were subsequently fragmented on ice using a sonicator (Diagenode Biorupter^TM^ UCD-300 bath sonicator set to high, 30s sonication followed by a delay period of 30s, 10 min total) to form PFFs.

### Assessment of fibrillar material

TEM and PAGE were used to characterize fibrillar αS (Fig EV1). For TEM, 10 μL of aggregated protein samples (from both before and after sonication) were incubated on TEM grids (400-mesh Formvar carbon coated copper, Electron Microscopy Sciences) for 1–2 minutes. Sample solution was wicked with filter paper and grid was washed with water to remove any excess material and improve background contrast. Grids were then incubated with 1% (w/v) aqueous uranyl acetate (UA) solution (10 μL) for 30–60 s. Excess UA was wicked away with filter paper and grids were air dried. TEM images were collected using a JOEL JEM 1011 TEM (operating voltage 100 kV) equipped with a charge-coupled device camera (ORIUS 832. 10W; Gatan). For PAGE, aggregated protein solutions were centrifuged to pellet the aggregated material. The upernatant was removed and pellet was resuspended in the starting volume of buffer. Both supernatant and resuspended pellet (20 μL) were loaded on a 4-12% polyacrylamide gel. Gels were imaged using a Typhoon FLA7000 gel imager (GE Life Sciences) using Coomassie stain mode.

### Cell culture

Human neuroblastoma (SH-SY5Y) and human embryonic kidney 293T (HEK) cells were grown at 37 °C under a humidified atmosphere of 5% CO_2_. The SH-SY5Y cells were cultured in Dulbecco’s Modified Eagle’s Medium (DMEM) plus 10% fetal bovine serum, 50 U/ml penicillin and 50 μg/ml streptomycin. The HEK cells were cultured in DMEM supplemented with 10% FBS, 2mM L-glutamine and 100 units/ml penicillin-streptomycin.

Cells were passaged upon reaching ~95% confluence (0.05% Trypsin-EDTA, Life Technologies), propagated and/or used in experiments. Cells used in experiments were pelleted and resuspended in fresh media lacking Trypsin-EDTA.

### Primary Neuronal Culture

Primary neuronal cultures were obtained from the Neurons-R-Us facility at the University of Pennsylvania. They were prepared from E15-E17 embryos of CD1 mice. All procedures were performed according to the NIH Guide for the Care and Use of Experimental Animals and were approved by the University of Pennsylvania Institutional Animal Care and Use Committee (IACUC). Dissociated hippocampal neurons were plated onto sterile, poly-D-lysine coated on IBIDI chambers at 200,000 cells/coverslip for live cell imaging and were allowed to mature for 5 days in complete neuronal medium (Neurobasal without phenol red (Thermo Fisher), 5% B27 supplement (Thermo Fisher). Medium was partially exchanged every 3-4 days.

### Giant Plasma Membrane Vesicles

Giant plasma membrane vesicles (GPMVs) are blebs obtained directly form the cell plasma membrane that contain lipid bilayers and the embedded membrane proteins, but lack the other biological components of the cell (Bauer et al, 2009). GPMVs were isolated from SH-SY5Y and HEK cells according to established methods (Sezgin et al, 2012). Briefly, cells were plated in 25 cm^2^ culture flasks and cultured for 48 h, washed with GPMV buffer (10 mM HEPES, 150 mM NaCl, 2 mM CaCl_2_, pH 7.4) twice and then exposed to 25 mM formaldehyde and 2 mM DTT for 2 h to induce blebbing. To reduce the content of DTT, GPMVs were dialyzed in GPMV buffer prior to use in experiments. GPMVs were also created using *N*-ethylmaleimide as the blebbing reagent, with comparable results. The phospholipid content of final material was measured by total phosphate assay.

### Phosphate assay

Lipid concentrations for GPMV preparations were estimated by measuring total phosphate, assuming that all measured phosphate is from phospholipids, and that all lipids are phospholipids. This is a practical assumption designed to ensure reproducibility.

### Enzymatic cleavage of carbohydrates

For cleavage of carbohydrates from GPMVs, endoglycosidases were added to the GPMVs in GPMV buffer at concentrations recommended by the manufacturers (PNGase F - 5,000 units/ml; Endo H - 2500 units; Heparinase I/III - 2500 units) and incubated at 37°C for 6 h. PNGase F was tagged with a chitin binding domain (Remove-IT PNGase F, New England Biolabs, MA, USA). For the images shown in this manuscript, the enzyme was removed by incubation of GPMVs with 50 μl of chitin binding magnetic beads. However, control experiments were conducted without removal of PNGase F and found to be comparable. Cleavage of N-linked glycans by PNGase F was confirmed by comparing images of GPMVs +/- PNGase F treatment after incubation with 50 nM concanavalin A (conA), a lectin which binds to binding to α-D-mannose and α-D-glucose moieties, or wheat germ agglutinin, a lectin which binds to N-acetyl-D-glucosamine and sialic acid. A significant decrease in the amount of both proteins is observed in these images (Fig EV6B).

For cleavage of carbohydrates from cells, cells were first plated for 42 h. After 42 h, media was removed from cells and replaced with FBS-free media complimented with the endoglycosidase (PNGase F – 5,000 units/ml; Endo H – 2500 units; heparinase I/II – 2500 units). The cells were incubated at 37°C under a humidified atmosphere of 5% CO_2_ for an additional 6 h. The media was then removed and replaced with cell growth media prior to the addition of αS. Cleavage of N-linked glycans from cells was confirmed by comparing images +/- PNGase F treatment after incubation with 50 nM conA-AL488, showing a significant reduction in the amount of conA bound (Fig EV5D).

To ensure that PNGase F is removed from GPMVs or cells after incubation (and therefore does not remain associated with either, blocking potential αS binding sites), we compared the amount of PNGase F added to either GPMVs or cells with that removed after incubation. PNGase F (5,000 units/ml) containing a chitin domain was added to chambers containing either GPMVs or cells and incubated at 37°C for 6 h, as for the experiments described above. After incubation, the buffer or media containing PNGase F was removed incubated with chitin magnetic beads to isolate and concentrate the enzyme. Blank chambers containing only PNGase F in buffer or media were subjected to the same treatment. 20 μL of each sample was run on a 4-12% polyacrylamide gels and stained with Coomassie blue. Gels were imaged using a Typhoon FLA7000 gel imager (GE Life Sciences) using Coomassie stain mode. The gels indicate that essentially all of the enzyme is removed (Fig EV9A).

### Quantification of carbohydrates

Concentrations of carbohydrates isolated from GPMVs or cells were quantified by using the Total Carbohydrate Quantification Assay Kit (Abcam, MA, USA) following the manufacturer’s instructions. Briefly, the carbohydrates are first hydrolyzed to monomer sugar units and then converted to furfural or hydrofurfural. These compounds are converted to chromogens, which can be detected by absorbance at 490 nm. Glucose was used to generate a standard curve for calculated of the total carbohydrate concentration of the samples.

### αS-captured carbohydrate pulldown assay

Carbohydrates cleaved and isolated from GPMVs (50 μg) were incubated with 100_□μM αS in 100_□μL 10 mM HEPES, 150 mM NaCl, 2 mM CaCl_2_, pH 7.4 for 1□h at room temperature. Binding reaction mixes were transferred to Amicon Ultra (3000 Da MWCO) centrifugal concentration devices that had been washed with 500□L deionized water. The concentrators were centrifuged for 5□min at 4200 rpm and the filtrates collected. The flow-through contains carbohydrates that did not bind αS and thus were not retained in the chamber of the concentrator. The amount of carbohydrate in the flow-through was quantified by the total carbohydrate assay as described above. The fraction of carbohydrate bound and retained by αS (*C_captured_*) was calculated relative to the starting concentration (50 μg) of the carbohydrate mixture, using;

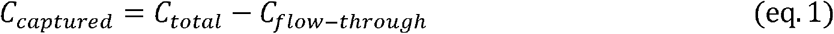

Where *C_total_* corresponds to the absorption of the starting stock concentration of carbohydrates used and *C_flow-through_* is the absorption of the flow-through.

Samples containing αS with no glycans or conA with 50 μg glycans were used as negative and positive controls, respectively. As expected, no signal at 490 nm was detected for the αS-only sample; the conA results are shown in Fig EV7F. The absorbance of flow-through at 280 nm was measured, confirming that all protein was retained in the concentrator and could not be detected in the flow-through.

### Cell imaging and analysis

All cell imaging was carried out by confocal fluorescence microscopy using an Olympus FV3000 scanning system configured on a IX83 inverted microscope platform with a 60x Plan-Apo/1.1-NA water-immersion objective with DIC capability (Olympus, Tokyo, Japan). For all experiments the gain setting for the blue channel was kept constant from sample to sample (excitation 488 nm, emission BP 500-540 nm). For detection of αS-AL594, the green channel was used (excitation 561 nm, emission BP 570-620 nm). Images were obtained in 8-well ibidi chambers (μ-Slide, 8-well glass bottom, ibidi GmbH, Germany) coated with Poly-D-lysine and were seeded with 20000-25000 cells/well. Cells were cultured for 48 h after passage before beginning experiments. For cellular uptake experiments, 200 nM αS-AL488 was incubated with cells for 0 to 24 h before acquiring images. For experiments using deglycosylated cells, cells were pretreated with the tested endoglycosidase for 6 h as described above prior to addition of protein. For colocalization with lysosomes, cells were treated with 75 nM Lysotracker Deep Red (Life Technologies) for 1 h prior to imaging. For all experiments, the gain setting for each channel was kept constant from sample to sample.

Image acquisition and processing were performed with the software accompanying the FV3000 microscope and Image J software (Schneider et al, 2012). For SH-SY5Y cells, internalized αS was quantified either by analysis of the punctate structures in the cells or by the total cellular fluorescence; for primary neurons, internalized αS was quantified by total cellular fluorescence.

For total cellular fluorescence, the integrated fluorescence intensity of the cells is calculated and reported. Cellular puncta were analyzed using the Image J particle analysis plug-in. This algorithm detects puncta through a user-defined threshold and counts the number of puncta that meet or exceed the threshold. The threshold was initially defined by manual identification and averaging of a subset of puncta. Colocalization with lysosomes was computed by obtaining a Pearson coefficient using the ImageJ plugin for co-localization (Coloc_2).

### Endocytosis inhibition

For inhibition of exocytosis experiments, monomer αS-AL488 (200 nM) or PFFs (200 nM in monomer units, 20:1 αS:αS-AL488) was initially incubated with cells for 30 minutes at 4°C. Control cells were moved to 37°C while the endocytosis inhibited cells were incubated at 4°C for another 4 h before acquiring images. For all experiments, the gain setting for each channel was kept constant from sample to sample. Image acquisition and processing were achieved using the software accompanying the FV3000 microscope and Image J software (Schneider et al, 2012).

### GPMV imaging and analysis

All GPMV images were carried using a PicoQuant MicroTime 200 time-resolved fluorescence system based on an inverted Olympus IX73 microscope (Olympus, Tokyo, Japan) with a 60x Plan-Apo/1.4-NA water-immersion objective using a 482 nm excitation laser and a frame size of 512 × 512 pixels. Images acquired with this instrument were in lifetime mode but were integrated to obtain intensity based images comparable to typical confocal images. This instrument has the advantage of very sensitive avalanche photodiode detectors (SPADs) that are capable at detecting nM concentrations of protein. Fluorescence intensities were analyzed by via the lifetime mode (both intensity and FRET images) using SymPhoTime 64 (PicoQuant). The intensity of images was then adjusted on ImageJ analysis program (Schneider et al, 2012).

For FLIM-FRET experiments, measurements were made of donor-only and donor-acceptor labeled proteins. SPAD signals were processed with the TimeHarp 300 photon counting board and analyzed with the SymPhoTime 64 software (PicoQuant) taking into account the instrument response function to allow consideration of short lifetime components with a high accuracy. FLIM images were acquired for 180 s, with a pixel integration time of 40 μs per pixel and an average photon count rate of 10000-30000 counts per second. Regions of interest of the GPMV membrane were selected from FLIM images and fluorescent lifetimes were obtained from TCSPC decay curves fitted by an exponential equation using the SymPhoTime 64 software. Fitting of the fluorescence images was then performed pixel wise with a single exponential model on all pixels above an intensity threshold of 200 photons. By characterizing donor lifetime in the absence and presence of acceptor, FRET efficiency (ET_eff_) can be calculated from:

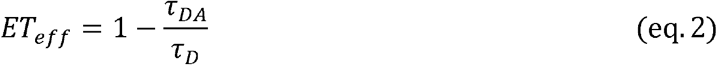

where τ_DA_ and τD and are the donor excited state lifetime in the presence and absence of acceptor. Six FLIM images were recorded for each of three biological repeats per condition. The histograms shown in Fig 5 represents ET_eff_ values for selected pixels from equatorial sections of the GPMVs as indicated on the images to their left. The histograms were fit with a Gaussian function to extract the mean ET_eff_.

Images were obtained in 8-well NUNC chambers (Thermo Scientific, Rochester, NY, USA) coated with Poly-D-lysine, containing 250 μl of GMPV at 5 μM phospholipid concentration and 80 nM of αS labeled with Alexa 488. For all experiments using these chambers, the chambers were passivated by polylysine-conjugated PEG treatment to prevent any nonspecific absorption to the chamber surfaces (Middleton & Rhoades, 2010).

### Fluorescence correlation spectroscopy (FCS)

Fluorescence correlation spectroscopy (FCS) measurements were carried out on a lab-built instrument, as described previously (Trexler & Rhoades, 2009). A 488-nm-wavelength laser was adjusted to 5 μW prior to entering the microscope. Fluorescence emission was collected through the objective and separated from laser excitation using a Z488RDC Long-Pass Dichroic and an HQ600/200M Band-Pass Filter (Chroma). The fluorescence emission was focused into the aperture of a 50-μm-diameter optical aperture fiber (OzOptics) directly coupled to an avalanche photodiode. A digital correlator (Flex03LQ-12; Correlator.com) was used to generate the autocorrelation curves. For each experiment, 30 autocorrelation curves of 10 s each were acquired and averaged together to obtain statistical variations. These average autocorrelation curves were then fit to a function for single fluorescent particles undergoing Brownian motion in a 3D Gaussian volume weighted by the inverse square of the SD:

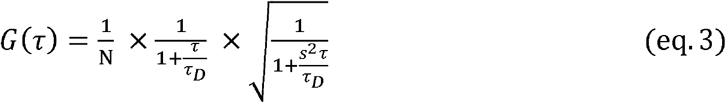

where G(τ) is the autocorrelation function for translational diffusion as a function of time τ, N is the average number of fluorescent molecules in the laser focal volume, and τ_D_ is the translational diffusion time of the particles. The structure factor, s, is the ratio of the radial to axial dimensions of the focal volume and was obtained from a calibration procedure using Alexa 488 hydrazide dye solutions (s = 0.2) and fixed for all subsequent fitting. Several datasets of autocorrelation curves obtained in the presence of carbohydrates were analyzed considering a single component system. For carbohydrate binding, both the diffusion time, τ_D_, (Fig 6A, 6B, EV7A, EV7C and EV7D) and the number of molecules, N, (Fig 4C, 4E and EV7B) were analyzed.

### GPMV binding and cellular uptake by FCS

FCS was used to monitor αS both binding to GPMVs and uptake by cells. FCS measurements were made on the instrument described above. For GPMV binding, experiments were performed in 8-well NUNC chambers (Thermo Scientific, Rochester, NY, USA) containing 250 μl of GMPV at 5 μM phospholipid concentration. The laser focal volume was located to a height of 100 μm from the bottom surface of the wells, above all GMPVs. αS labeled with Alexa 488 (80 nM) was added to wells with GPMVs and the autocorrelation measurements were taken immediately and after 60 minutes. Each curve was integrated for 30 s, repeated 10 times and data was analyzed as described above. Binding was quantified by measuring the change in the number of molecules, N, of fluorescently labeled αS present in the solution surrounding the GMPVs relative to the starting concentration. There was no change in the diffusion time of the αS that remained in the cell media over this time course, evidence that protein is stable and the fluorophore intact (Fig EV9B).

For cell uptake experiments, cells were plated in phenol red-free medium in 8-well ibidi chambers (μ-Slide, 8-well glass bottom, ibidi GmbH, Germany) coated with Poly-D-lysine wells and cultured for 48 h prior to measurements. Control wells contained the same volume of media without cells; these control for non-specific adsorption of protein to the well surfaces. αS-AL488 (200 nM) was added to the wells at the start of the experiment, and the autocorrelation curves were taken in the medium well above the cells, at a height of 100 μm from the bottom surface of the wells. The curves were collected at regular intervals, over a period ranging from 2 to 24 h, and each autocorrelation curve integrated for 30 s, repeated 10 times. Data were analyzed as described above. Cellular uptake was assessed by measuring the change in the number of molecules of fluorescently labeled αS present in the media surrounding the cells relative to the starting concentration. In between measurements, the chambers were returned to the incubators to maintain the temperature in the wells at 37°C. As a negative control for both the GPMV binding and cell uptake studies, GPMVs or cells were incubated with 80 nM eGFP. There was no evidence of eGFP binding to the GPMVs nor of internalization by SH-SY5Y cells (Fig EV9C).

### Cellular uptake by PAGE

As an orthogonal approach to the FCS and imaging approached described above, uptake of both monomer and PFF αS were quantified by PAGE. SH-SY5Y cells were incubated with 200 nM αS, as described above for FCS uptake experiments. After incubation for the desired time (1, 3, 5, 8, 12, or 24 h) the media was removed from cells and stored at 4°C. After all samples were collected, 20 μl of each sample were run on a 4-12% polyacrylamide gel.

To quantify the amount of protein internalized, an identical set of experiments was carried out, with modifications as described in the following. For each time point, transferrin-AL488 (100 nM) was added 30 min prior to the due time of the time point to serve as a loading control. At the desired time points, the cells were detached from the wells by 0.05% Trypsin-EDTA (Life Technologies), pelleted and lysed in 250 μl RIPA lysis buffer (Thermo Fisher). Cell lysates (20 μl of stock) were run on PAGE gels. Cell lysates (20 μl) were run on 4-12% polyacrylamide g^els^.

Gels of both extracellular and internalized αS were imaged using a Typhoon FLA7000 gel imager (GE Life Sciences) using fluorescent imaging mode to detect αS-AL488 or transferrin-AL488. Image J was used to quantify the bands.

### Statistical analysis

Data are expressed as the mean □±□ SD and were examined by a one-way analysis of variance (n□=□3). More than three experiments were performed and similar results were obtained. P values less than 0.05 were considered to be significant.

### Trypan Blue Quenching

Trypan blue solution 0.4% (Thermo Fisher) and fresh neurobasal without phenol red, B27, or antibiotic supplementation were equilibrated at 37°C. A 10X dilution of trypan blue was prepared freshly in the warmed neurobasal media. The trypan blue solution was then added to cells dropwise and incubated at 37°C for an hour prior to imaging. Trypan Blue quenching was performed for all imaging experiments performed using PFFs and when monomer protein was introduced in experiments utilizing primary neurons. Trypan Blue quenching was used eliminate signal from extracellular material allowing the fluorescence quantification of intracellular fluorescence signal.

### Propagation of amyloid in primary neurons

Primary wild type mouse hippocampal neurons (obtained as described above) were grown for 6 days on round coverslips prior to addition of αS_acetyl_ or αS_un_ PFFs (100 nM final PFF concentration). Cells were fixed with 4% (wt/vol) paraformaldehyde and co-stained with antibodies specific to αS phosphorylated at serine 129 (rabbit monoclonal phospho 129, 1:250 dilution) and neuronol tubulin (mouse monoclonal anti-β-III tubulin, 1:100 dilution) after 3-, 7- and 10-days incubation with PFFs. Primary antibodies were visualized by secondary staining with Alexa Fluor 488 donkey anti-rabbit IgG (Invitrogen) and Alexa Fluor 647 goat anti-mouse IgG (Invitrogen) (1:1000 dilution).

### Transfection of HEK cells

HEK cells were transfected with plasmid encoding LAG3-GFP or Neurexin 1 β-GFP by Lipofectamine 3000, following manufacture’s directions. The media was removed from cells 48 h after transfection and replaced with FBS-free media complimented with PNGaseF (5,000 units/ml) for experiments that required PNGaseF treatment. The cells were then incubated at 37°C under a humidified atmosphere of 5% CO2 for an additional 6 h. The media was then removed and replaced with cell growth media prior to the addition of αS. Cells were incubated with monomer αS labeled with Alexa 594 (final concentration 200 nM) or PFF αS (final concentration 200 nM monomer units, 1:20 labeled:unlabeled) for 12 h prior to imaging.

### Flow cytometry

To quantify LAG3-GFP and Neurexin 1 β-GFP expression levels, HEK cells were plated and transfected as above. Following 48 h, cells were detached using 0.05% Trypsin-EDTA, centrifuged and washed with PBS + 2mM EDTA and 2% BSA. The cells were then treated with the Zombie Yellow fixable cell viability kit (BioLegend) at room temperature in PBS + 2mM EDTA for 20 mins and then fixed at 4 □C using Cytofix for 20 mins. Cells were then placed in PBS + 2mM EDTA and 2% BSA. Data were collected on an LSR II flow cytometer (BD Biosciences) and post-collection data were analyzed using FlowJo (Treestar).

### Nuclear Magnetic Resonance

^1^H-^15^N HSQC NMR titrations were carried out at 25°C using Varian 600 MHz or Agilent 800 MHz spectrometers equipped with room temperature probes. A uniformly labeled ^15^N-αS solution was added to either N-linked glycans obtained by PNGaseF cleavage from SH-SY5Y cells or commercially available mono- or trisaccharides at the concentrations indicated in Fig 6C and EV7E in GPMV buffer (10 mM HEPES, 150 mM NaCl, 2 mM CaCl2, pH 7.4, 10% D_2_O; final concentration of ^15^N-αS was 350 μM). HSQC spectra were collected with VnmrJ software using built-in pulse sequence including WATERGATE solvent suppression and analyzed with Mnova software suite (Mestrelab). Standard parameters for zero-filling, apodization and baseline correction were applied. ^1^H chemical shifts were referenced using water resonance and ^15^N chemical shifts were referenced indirectly based on gyromagnetic ratios of respective nuclei. Previously assigned αS backbone resonances were used (Kang et al, 2011).

Binding to the PNGaseF derived glycans results in non-uniform peak intensity increases throughout the sequence of αS. For analysis, 10 peaks showing large increases and 10 peaks showing small or no increases were selected to roughly cover the entire sequence. The same residues were used for each analyzed dataset. For each set of peaks, the relative magnitude of increase (as compared to αS in solution without glycans) expressed as a percentage was calculated and averaged. A difference between two numbers is reported in Fig 6C of the manuscript. The same calculation was carried out for the mono- and tri-saccharides, which do not bind to αS, shown in Fig. EV7E.

## Supporting information

## Acknowledgments

This research was supported by NIH R01 NS079955 and the Michael J. Fox Foundation (to E.R.). We thank T. Baumgart for use of his confocal microscope, V. M.-Y. Lee for use of her sonicator, M. Maronski of the Neurons R Us Culture Service Center at Penn Medicine Translational Neuroscience Center (PTNC) at the University of Pennsylvania for hippocampal neuron preparation, J. Wang for advice on the neuron propagation assay, J. Doerner for the use of the flow cytometry and R. Jin for providing the vesicle schematic adapted for Fig 5. All data are available from authors by request.

## Author contributions

M.B. and E.R. designed and conceived the experiments. M.B. performed all measurements on GPMVs, cells, and isolated glycans. S.P.W performed the NMR experiments in consultation with A.D.M. All figures and text were prepared by M.B. and E.R.

## Expanded View Figure legends

**Expanded View Figure 1. Characterization of monomer and PFF αS.**

A MALDI-TOF mass spectrometry was used to confirm the presence of the N-terminal acetyl group as well as the purity of the samples for both unlabeled and Alexa 488 labeled αS. The expected masses for αS_acetyl_^E130C^ and αS_acetyl_^E130C^-AL488 are 14,576 and 15174, respectively; for αS_un_^E130C^ and αS_un_^E130C^-AL488 are 14434 and 15132, respectively; the reported values are within expected accuracy for MALDI-TOF.

B The fibril morphology before and after sonication was examined by TEM at 100,000x magnification. Scale bar=200 μm.

C PAGE analysis at the end of the aggregation assay indicates that the majority of the material (αS_acetyl_) is fibrillar. 1= molecular weight standards; 2 = pellet; 3 = supernatant

**Expanded View Figure 2. Time dependent endocytosis of αS_acetyl_ monomer and PFFs.**

A Time dependent uptake of αS_acetyl_ (green) monomer or PFFs by untreated or PNGase F treated SH-SY5Y cells. Incubation time indicated above each image. Cells were stained with LysoTracker Deep Red (purple) prior to imaging.

B Image overlap statistics for (A) of LysoTracker and αS_acetyl_ monomer and PFFs at the indicated incubation time. Colocalization was analyzed with the Pearson correlation coefficient. A larger coefficient reflects more overlap between αS_acetyl_-AL488 puncta and LysoTracker puncta (endosomes).

Data information: Correlation coefficient was computed using the ImageJ plugin for colocalization (n=100 cells, 3 independent experiments, **P<0.01, ***P<0.001 and ****P<0.0001 by the Student’s T-Test. Scale bars=20 μm.

**Expanded View Figure 3. Quantification of cell uptake measured by PAGE analysis.**

Uptake of αS_acetyl_ monomer (200 nM αS_acetyl_-AL488) or PFFs (200 nM in monomer units, 20:1 αS_acetyl_: αS_acetyl_-AL488) by SH-SY5Y cells as measured by PAGE analysis with fluorescence imaging of the gels (to detect only αS_acetyl_-AL488). (Upper) Gels show αS_acetyl_ remaining in the media at the timepoints indicated above the gels. Uptake is measured by quantifying the decrease of αS_acetyl_-AL488 in the media as a function of time. Quantification of the gels is shown as the scatter plot for monomer and PFF αS_acetyl_ +/- PNGase F treatment. The measurements are analogous to the FCS measurements shown in Fig 2C in the main manuscript and the results of both approaches are comparable. (Lower) Gels show αS_acetyl_ internalized by cells at the time points indicated above the gels. Uptake is measured by quantifying the amount of αS_acetyl_-AL488 from lysed cells as a function of time. Transferrin-AL488, which exhibits very rapid uptake kinetics (Fig EV4C), was added to cells for 30 minutes prior to lysis, to be used as loading control. The scatter plot compares the amount of internalized monomer and PFF αS_acetyl_, and the bar plot compares the amount of both forms internalized +/- PNGase F treatment. Quantification of gel band intensity was computed using ImageJ. These measurements are analogous to the image analysis shown in Fig 2C and the results from both approaches are comparable.

Data information: For each experiment, 3 independent measurements were made.

**Expanded View Figure 4. Clathrin dependent endocytosis.**

A Inhibition of endocytosis monitored by uptake of αS_acetyl_ monomer or PFFs at 4°C. Images are shown of uptake of protein at 37°C and 4°C for each condition. Protein was added to the cells followed by incubation at 4°C for 30 mins. The controls were then moved to 37°C incubator. The two groups of cells were incubated in the respective temperatures for an additional 4 h.

B HEK cells incubated with αS _acetyl_ monomer or PFFs for 12 h.

C Rates of clathrin-dependent endocytosis of SH-SH5Y and HEK cells compared by loss of 100 nM transferrin-AL488 from extracellular medium of cells as measured by FCS.

Data information: All αS_acetyl_ uptake measurements used 200 nM αS-AL488 monomer or PFF (concentration in monomer units, 20:1 αS:αS-AL488) Scale bar=20μm.

**Expanded View Figure 5. Treatment of cells with endoglycosidases.**

A Heparinase treatment of SH-SY5Y cells disrupts uptake of αS_un_ PFF (upper) but not of αS_acetyl_ PFF (lower). Images made after 12 h of incubation of SH-SY5Y cells with 200 nM PFF αS_acetyl_-AL488 or αS_un_-AL488 (concentration in monomer units, 20:1 αS:αS-AL488) following treatment with Heparinase.

B Heparinase and Endo H treatment of SH-SY5Y cells does not disrupt uptake of monomer αS_acetyl_. Images were made after 12 h of incubation of SH-SY5Y cells with monomer αS_acetyl_-AL488 (200 nM) following treatment with Endo H or Heparinase. Quantification of monomer αS_acetyl_ uptake by SH-SY5Y cells treated with Endo H or Heparinase shown relative to PNGase F treated or untreated cells (Fig 2E). Numbers of puncta were computed using the ImageJ plugin for particle analysis (n=100 cells, 3 independent experiments, **P<0.01, ***P<0.001 and ****P<0.0001 by the Student’s T-Test.

C 50 nM conA-AL488 incubated with SH-SY5Y cells +/- PNGase F treatment. A significant reduction in the amount of conA-AL488 is observed in cells treated with PNGase F.

D (Upper) Uptake of transferrin and αS_acetyl_ by PNGase F treated SH-SY5Y cells. Internalization of transferrin-AL488 is not impacted by this treatment, indicating that clathrin-mediated endocytic pathways are functional. Uptake of αS_acetyl_-AL594 is significantly reduced. (Lower) As with the SH-SY5Y cells, uptake of transferrin by primary neurons from embryonic mouse hippocampus is not impacted by PNGase F treatment, indicating that clathrin-mediated endocytic pathways are functional.

E Colorimetric measure of toxicity following the incubation of SH-SY5Y cells with 200 nM αS_acetyl_ monomer or PFFs (concentration in monomer units) for times indicated on plots. Data are expressed relative to vehicle-only addition prepared on the same well-plates. Each histogram bar is the average of eight, on-plate repeats across each of three independently performed replicates (n = 24).

**Expanded View Figure 6. αS binding to and clustering of SH-SY5Y GPMVs is dependent on N-linked glycans.**

A Representative images of SH-SY5Y GPMVs incubated with 100 nM αS_acetyl_-AL488 and 80 μM of unlabeled αS_acetyl_.

B Representative images of PNGase F treated SH-SY5Y GPMVs incubated with 100 nM αS-AL488 and 80 μM of unlabeled αS.

C Representative images of Endo H and Heparinase treated SH-SY5Y GPMVs incubated with 100 nM αS_acetyl_-AL488 and 80 μM of unlabeled αS_acetyl_.

D Representative images of HEK GPMVs incubated with 100 nM αS_acetyl_-AL488 and varying concentrations of unlabeled αS_acetyl_ (indicated).

E GPMVs incubated with 50 nM conA-AL488 or 50 nM wheat germ agglutinin-AL488, +/-PGNase F treatment (upper/lower).

F GPMVs incubated with 100 nM αS_acetyl_-AL594 and 80 μM of unlabeled αS_acetyl_ prior to addition of 50 nM conA-AL488.

G GPMVs incubated with 80 μM of unlabeled αS_acetyl_ prior to the addition of 50 nM conA-AL488.

Data information: For all experiments, GPMVs equivalent to 5 μM total lipid as measured by the phosphate assay were used. Scale bars=20 μm.

**Expanded View Figure 7. αS binding to isolated glycans.**

A Averaged autocorrelation curves (30 curves of 10 seconds each) and fits to Equation 3 for αS_acetyl_ in the presence and absence of PNGase F cleaved glycans.

B The number of αS_acetyl_ molecules, N, upon, titration with PNGase F cleaved glycans by FCS (same measurements as analyzed for diffusion time in Fig 6A). A decrease in N as a function of glycan concentration would reflect aggregation or oligomerization of the protein; this is not seen here.

C Diffusion time of 80 nM conA-AL488 as a function of increasing concentrations of PNGase F-derived glycans.

D Diffusion time of 80 nM αS_acetyl_-AL488 as a function of increasing concentrations of Endo H and Heparinase-derived glycans. Data for PNGase F derived glycans also shown for comparison (Fig6A).

E Average increase in peak intensity for strongly affected residues relative to weakly/unaffected residues from ^15^N-^1^H HSQC spectra of αS_acetyl_ in the presence of H-Trisaccharide (H-Tri) and N-acetylglycosamine (GlcNAc). The ratio of simple sugar to protein used in indicated.

F Quantification of PNGase F cleaved glycans bound to monomer and PFF forms of αS_acetyl_ using total carbohydrate assay (reported as absorption at 490 nm). For comparison, positive control conA is shown.

Data information: All results are relative to the initial glycan pool which is treated to the same filtration and quantification protocol as the samples. Details of the assay are described in the Materials and Methods.

**Expanded View Figure 8. Internalization of αS by HEK cells transfected with LAG3 or neurexin 1β.**

A HEK cells transfected with GFP-tagged neurexin 1β (upper) or GFP-tagged LAG3 (lower), imaged 48 h after transfection.

B Transfection efficiencies of GFP-tagged neurexin 1β (upper: ~21%) or GFP-tagged LAG3 (lower: ~16%) measured by flow cytometry. Lipofectamine-only sample was used as a GFP negative control used for gating purposes in the analysis.

C HEK cells transfected with GFP-tagged LAG3 as in (A) but with the addition of monomer αSun–AL594.

D HEK cells transfected with GFP-tagged neurexin 1β as in (A) but with the addition of monomer αS_un_-AL594.

Data information: All confocal imaging shown was conducted after 48 h transfection. For images (A, C and D) cells were incubated for an additional 12 h with 200 nM of αS_acetyl_ or αS_un_-AL594. For PFFs, 200 nM total concentration in monomer units (1:20 labeled:unlabeled). The data are presented as mean±S.D., n=3. Scale bars=20 μm.

**Expanded View Figure 9. PNGase F does not remain bound to cell membranes after incubation.**

A The amount of PNGase F before and after treatment of GPMVs and SH-SY5Y cells was examined by PAGE. The majority of the enzyme is recovered from the GPMV buffer or cell media, indicating that it does not remain bound to the membranes, potentially blocking αS_acetyl_ binding.

B αS_acetyl_ is stable during incubation with cell media during uptake studies. FCS was used to monitor the diffusion time of αS_acetyl_ in media as a function of time during incubation with SH-SY5Y cells. The diffusion time is stable over the 24 h period, indicating that the protein is not degraded nor is the fluorophore cleaved, both of which would be expected to result in a faster diffusion time.

C SH-SY5Y GPMVs and cells do not bind or uptake eGFP, respectively. eGFP was used as negative control. The addition of 80 nM GFP to GPMVs shows no evidence of binding. Likewise, there is no evidence of uptake of GFP by SH-SY5Y cells following 12 h of incubation. Scale bar=20μm.

