## Supplementary material for "Identification of N-linked glycans as specific mediators of neuronal uptake of acetylated α-Synuclein"

Data information: All αS_acetyl_ uptake measurements used 200 nM αS-AL488 monomer or PFF (concentration in monomer units, 20:1 αS:αS-AL488) Scale bar=20μm.

**Extended version 5. Treatment of cells with endoglycosidases.**

Extended version 1.

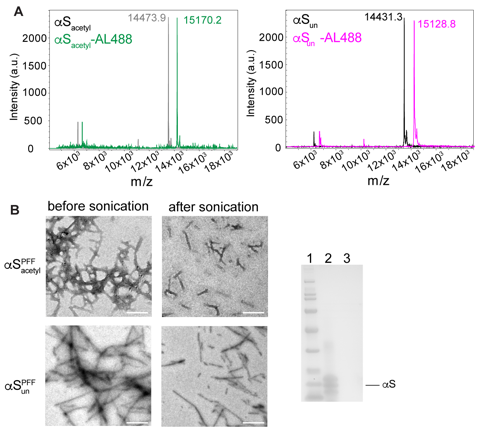

Extended version 2.

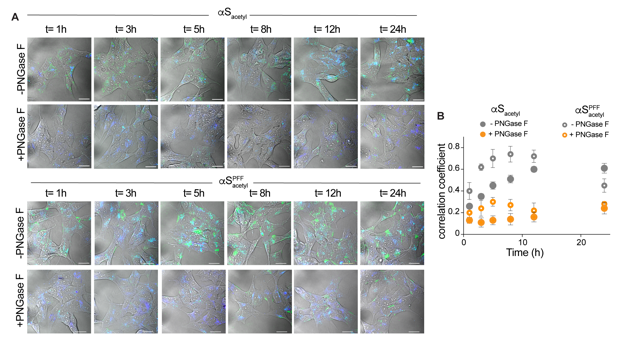

Extended version 3.

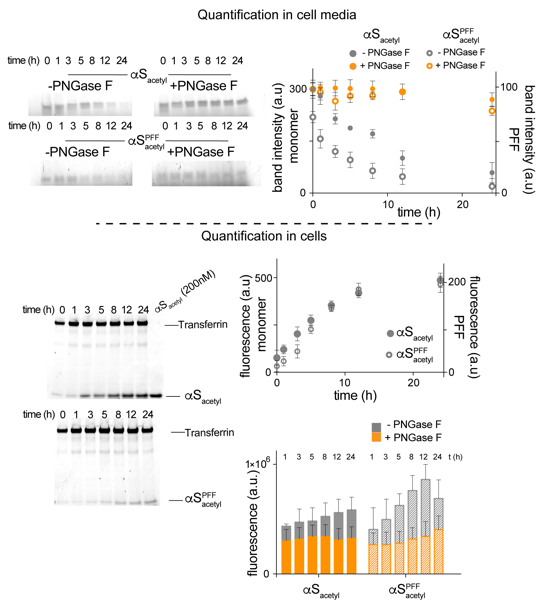

Extended version 4.

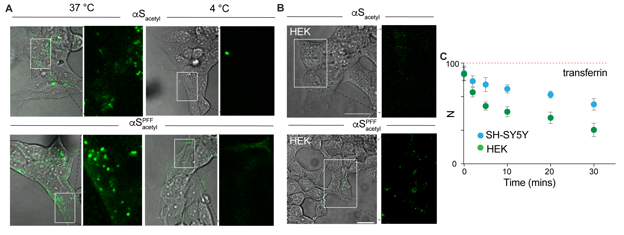

Extended version 5.

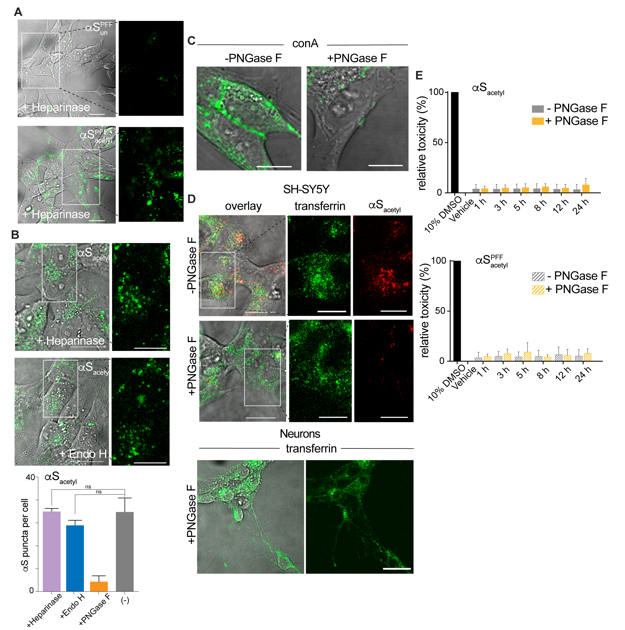

Extended version 6.

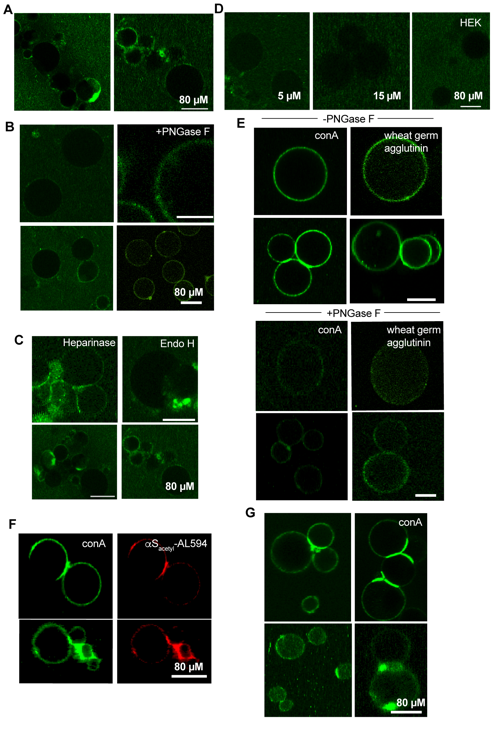

Extended version 7.

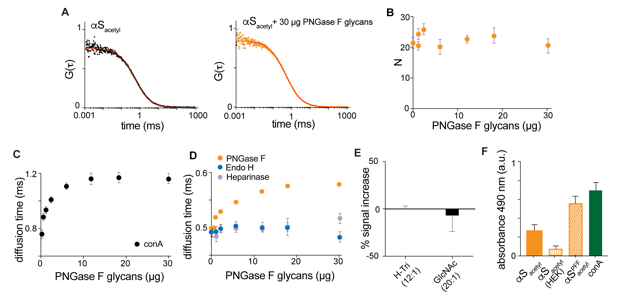

Extended version 8.

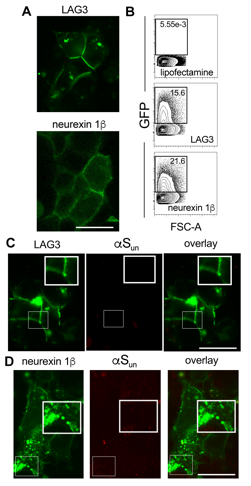

Extended version 9.

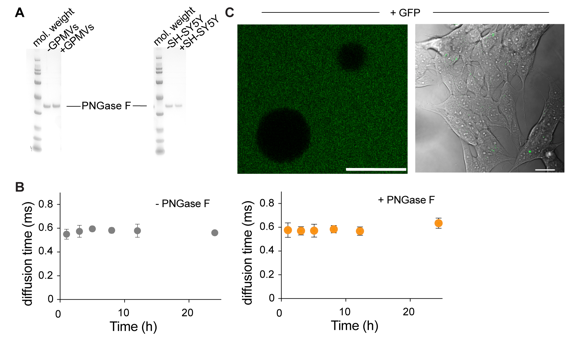
